# A novel tool for labeling intermediary proteins between two non-interacting proteins

**DOI:** 10.1101/2025.06.02.657518

**Authors:** Lin Xie, Hua Li, Lijuan Gao, Gangyun Wu, Wenxiu Ning

**Author notes:** Corresponding author (WN); (HL). These authors contributed equally to this work (Lin Xie, Hua Li, Lijuan Gao).

## Abstract

Decoding the complexities of signaling pathways is fundamental for deciphering the mechanisms underlying tissue development, homeostasis, and disease pathogenesis. Proximity labeling tools have been instrumental in identifying upstream or downstream effectors of specific proteins within signaling pathways. However, currently, there are no tools available to directly label and capture intermediary proteins that bridge two non-interacting proteins. Here, we developed STUPPIT (<u>S</u>plit-<u>Tu</u>rboID and <u>P</u>UP-IT based <u>P</u>rotein Identification <u>T</u>ool), a novel method combining split-TurboID and PUP-IT to biotinylate intermediary proteins of two non-interacting proteins through a two-step enzymatic reaction. Two approaches of STUPPIT were validated using three well-characterized protein triad, including YAP1/AMOT/β-actin, YAP1/LATS1/MOB1A, and β-catenin/α-catenin/β-actin. Combining STUPPIT and proteomics, we identified novel intermediary proteins including RNF20, ERC1, USP7 and TRIM33, which interact both with β-catenin and SMAD4, key components of the Wnt and BMP signaling pathways. In conclusion, STUPPIT represents a powerful tool for labeling and capturing intermediary proteins between non-interacting partners, offering new insights into protein-protein interactions and advancing signal transduction research.

## Introduction

Cellular signaling pathways depend on precisely regulated protein-protein interactions (PPIs) that form dynamic networks controlling fundamental biological processes. These interactions are not static but rather exhibit remarkable plasticity, enabling cells to adapt to microenvironments during tissue development, homeostasis, and stress responses. The identification of novel interacting partners of proteins of interest represents a crucial and central aspect of signaling pathway research (1-3).

Over time, versatile tools utilizing proximal enzymatic labeling have emerged for capturing proximal partners of known proteins, such as horseradish peroxidase (HRP) (4), ascorbate peroxidases (APEX2) (5), biotin ligases (BioID (6, 7), TurboID (8)), LIPSTIC (transpeptidase sortase A) (9) or its updated version EXCELL (10), and PUP-IT (pafA ligase (11)). These proximity labeling techniques allow for the identification of proteins that have transient and weak interactions within the proximity distance of an introduced labeling enzyme (3, 12, 13). The extended form of split proximity labeling enzymes, such as split-BioID, split-APEX2, and split-TurboID, facilitates the identification of proteins between two known interacting proteins (14-16). These proximity labeling tools have been instrumental in dissecting the upstream or downstream effectors of proteins of interest in signaling pathways, including Wnt, Hippo, Hedgehog, MAPK, and NF-κB pathways (17-22).

Despite significant advancements in the field, there remains a critical gap in our ability to directly label and capture intermediary proteins that bridge two associated but non-interacting proteins - a common scenario in signaling pathways. These intermediary proteins play a pivotal role in facilitating signal transduction and crosstalk, either by acting as adaptors to link non-interacting proteins or by competitively binding to them. For instance, in mechanotransduction, actin binds to the intermediary protein AMOT, thereby sequestering it from inhibiting YAP1 (23, 24). In adherens junctions, α-catenin serves as an intermediary protein that connects F-actin to junctional sites by interacting with β-catenin (25, 26). In the Hippo signaling pathway, MOB1 binds to the intermediary LATS1/2, which in turn facilitates the phosphorylation of YAP1 by LATS1/2, blocking its nuclear import and inhibiting its transcriptional programs (27-29). Moreover, in pathways such as Wnt and BMP, reciprocal interactions are essential for controlling stem cell activity during tissue homeostasis, as seen in the small intestine epithelium and hair follicles (30, 31). However, the mechanisms by which intermediary proteins might competitively bind to key components of these pathways to balance their activities remain poorly understood. A major reason for this gap in knowledge is the lack of tools to directly label intermediary proteins between two non-interacting but associated proteins.

To address this gap, we developed STUPPIT (<u>S</u>plit-<u>Tu</u>rboID and <u>P</u>UP-IT based <u>P</u>rotein Identification <u>T</u>ool), a novel proximity labeling tool specifically designed to label and capture intermediary proteins between two non-interacting proteins. STUPPIT integrates split-TurboID and PUP-IT methods to biotinylate intermediary proteins via a two-step enzymatic reaction. We utilized STUPPIT to verify known intermediary proteins for several pairs of proteins, including YAP1/AMOT/β-actin, YAP1/LATS1/MOB1A, and β-catenin/α-catenin/β-actin. All of these known intermediary proteins were successfully labeled and enriched using two different approaches of STUPPIT. Furthermore, by combining STUPPIT with proteomics, we identified novel intermediary proteins, such as RNF20, ERC1, USP7, and TRIM33, between β-catenin and SMAD4, which are key components of the Wnt and BMP signaling pathways. In conclusion, STUPPIT represents a powerful tool for labeling and capturing the intermediary proteins between two non-interacting proteins, thereby enhancing and expediting the investigation of protein-protein interactions in signal pathways.

## Results

### STUPPIT leverages split-TurboID and PUP-IT to enable labeling of the intermediary proteins between non-interacting proteins

To label the intermediary partner that bridges two associated but non-interacting proteins, we designed the STUPPIT tool by combining the concepts of split-TurboID and PUP-IT (11, 16).

The split-TurboID comprises two TurboID fragments (Tb(N), an N-terminal fragment, and Tb(C), a C-terminal fragment) split at the amino acid site L73/G74 (16). These split fragments can be attached to the two interacting proteins and brought together to reconstitute an active enzyme, allowing the labeling of proximal partners of two known interacting proteins (Fig 1A). On the other hand, PUP-IT is a proximity labeling method based on the prokaryotic enzyme pafA, which catalyzes the phosphorylation of the C-terminal Glu on Pup(E) and then conjugates the C-terminal Glu to the lysine residue on the target and proximal proteins (11) (Fig 1B).

**Fig 1.**
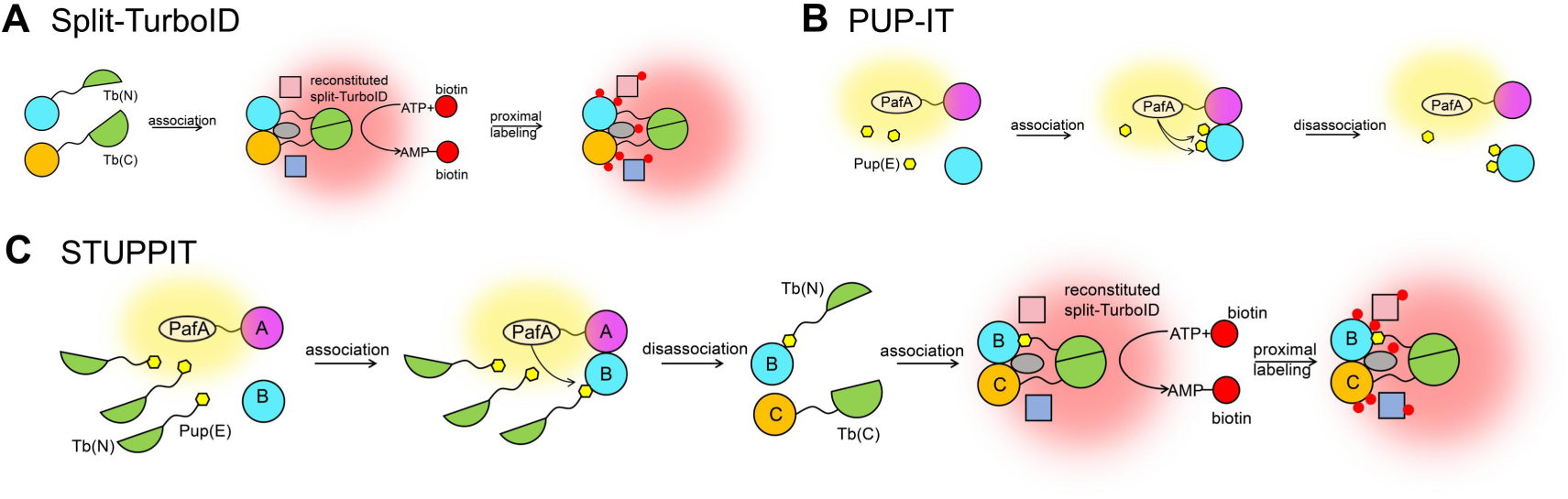
Design of STUPPIT for labeling and capturing intermediary proteins between two non-interacting proteins. (A-B) Schematic diagrams depicting the working principles of split-TurboID (A) and PUP-IT (B). (C) Schematic representation of how STUPPIT labels the intermediary proteins between two non-interacting proteins with biotin.

Taking advantage of the flexibility of modifying the N-terminus of Pup(E) with different tags without affecting its function, we opted to fuse the shorter fragment Tb(N) instead of the longer Tb(C) at the N-terminus of Pup(E) to create the Tb(N)-PupE substrate. The prey protein A is fused to pafA with a Myc tag. In the cellular environment with Tb(N)-PupE present, the neighboring interacting proteins of prey A, including the intermediary protein B, will be labeled with Tb(N)-PupE. As the intermediary protein B dissociates from Prey A and interacts with the Tb(C)-linked Prey C, the combined Tb(N) and Tb(C) will enzymatically reactivate to label the target proteins B and C along with their proximal proteins (Fig 1C). Hence, we named STUPPIT (<u>S</u>plit-<u>Tu</u>rboID and <u>P</u>UP-IT based <u>P</u>rotein Identification <u>T</u>ool) for this method designed to tag the intermediary connectors of two non-interacting proteins.

### Validating intermediary protein labeling efficiency using the all-in-plasmids approach I of STUPPIT

To evaluate the feasibility of the STUPPIT concept, we devised a workflow based on the all-in-plasmids approach. This method requires three constructs: the Prey A vector (PreyA-pafA-Myc), which links prey protein A with pafA-Myc; the control vector (HA-Tb(C)-IRES-3×Flag-Tb(N)-PupE), which co-expresses the Tb(C) control and the substrate Tb(N)-PupE; and the Prey C vector (HA-Tb(C)-PreyC-IRES-3×Flag-Tb(N)-PupE), which co-expresses Tb(C)-PreyC and the substrate Tb(N)-PupE. Each prey protein and substrate can be distinguished by different tags (Fig 2A).

**Fig 2.**
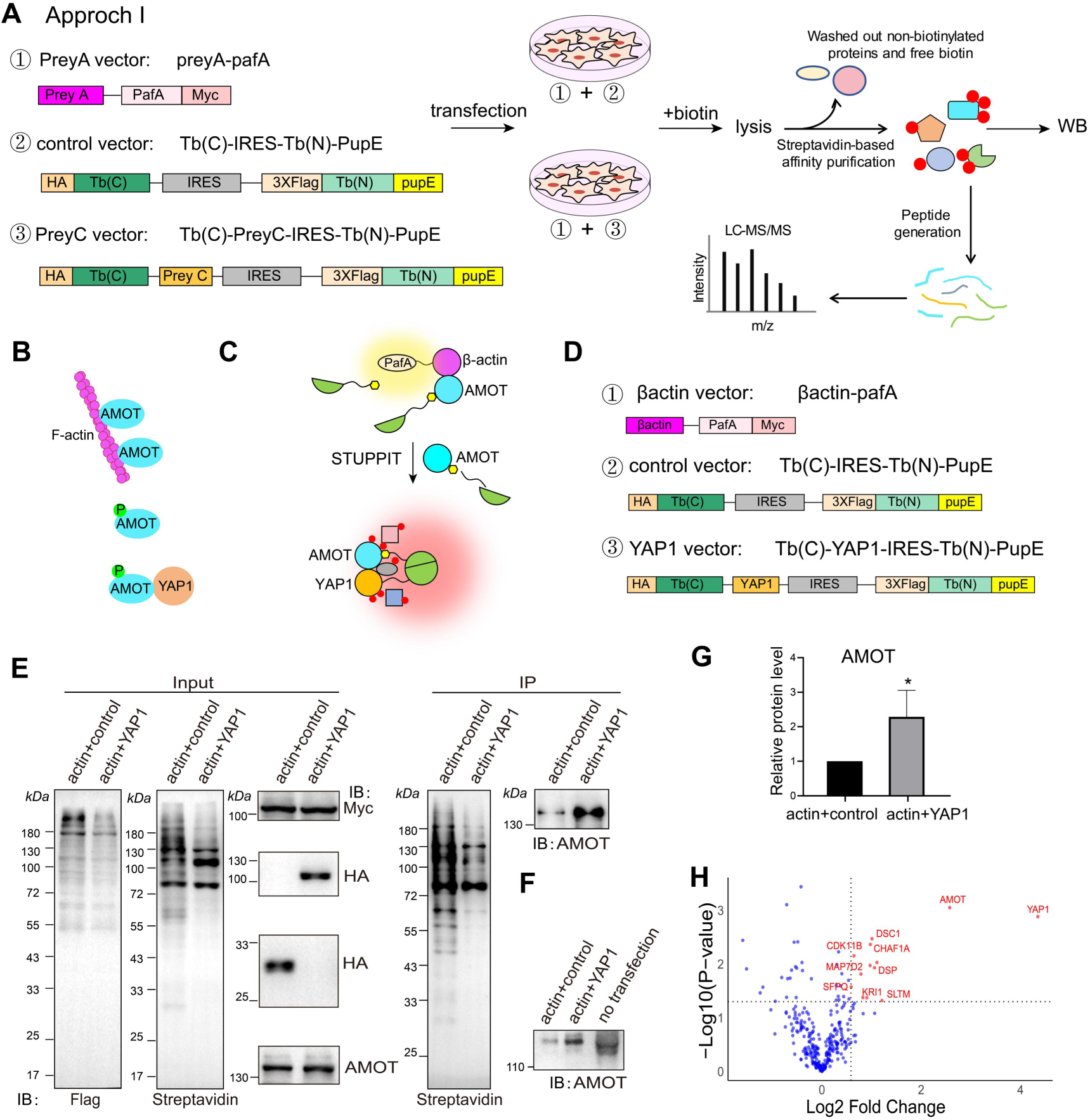
The all-in-plasmids Approach I of STUPPIT enables proximal labeling of the intermediary protein AMOT between YAP1 and β-actin. (A) Schematic diagram illustrating the all-in-plasmids approach I of STUPPIT for labeling and capturing intermediary proteins. (B) Illustration of AMOT as the intermediary protein between β-actin and YAP1. (C) Illustration of the identification of the intermediary protein AMOT between β-actin and YAP1 using approach I. (D) Schematic diagram outlining the plasmid construction required for approach I for the bait proteins of β-actin and YAP1. (E) Validation of the intermediary protein AMOT between YAP1 and β-actin captured by STUPPIT approach I through immunoblotting. (F) Immunoblotting showing the increased protein size of AMOT in STUPPIT. (G) Quantification of the immunoblotting intensity of AMOT between STUPPIT of β-actin&YAP1 compared to β-actin&control group. n=3 from 3 independent replicates. Data represents mean ± SD, n=3 for each group (three independent replicates), p-value = 0.0447, two-tailed unpaired t-test. (H) Volcano plots showing STUPPIT-enriched intermediary proteins in β-actin&YAP1 group compared to β-actin&control group using mass spectrometry analysis. Significantly enriched proteins with a 1.5 fold change are labeled as red dots. Proteins with unique peptides>3 in the β-actin&YAP1 group were selected.

To improve the efficiency of the STUPPIT system, we aimed to maintain only one variable, PreyC, in the experiment. By co-transfecting the preyA and preyC vectors, along with the preyA and control vectors, into cells and adding biotin, the intermediary proteins of prey A and C can be labeled with biotin. Subsequent steps involve utilizing streptavidin affinity immunoprecipitation and mass spectrometry proteomics to capture and identify the intermediary proteins (Fig 2A).

We first validated approach I of STUPPIT utilizing the well-known triad, F-actin/AMOT/YAP1. The F-actin/AMOT/YAP1 axis is a well-established interaction pathway during mechanotransduction, where β-actin competes with YAP1 for binding to AMOT, thereby inhibiting AMOT-mediated cytoplasmic retention of YAP1 (Fig 2B) (23, 24). Thus, AMOT serves as the intermediary protein between β-actin and YAP1, and should be enriched by STUPPIT (Fig 2C). Constructs were created for β-actin-pafA-Myc, the control vector (HA-Tb(C)-IRES-3×Flag-Tb(N)-PupE), and the YAP1 vector (HA-Tb(C)-YAP1-IRES-3×Flag-Tb(N)-PupE) as described previously (Fig 2D). The expression of these tagged proteins was validated through immunofluorescence staining and immunoblotting (Fig 2E and S1 Fig).

In the immunoblotting analysis, the 3×Flag-Tb(N)-PupE exhibited a single band when transfected alone into cells (S1C Fig). The gradient in anti-Flag blotting and the increase in AMOT protein size by STUPPIT indicated the efficient pafA-mediated catalysis of Tb(N)-Pup(E) on interacting proteins (Fig 2E-F). Furthermore, in the STUPPIT group of β-actin with YAP1, the streptavidin-recognized biotinylated proteins differed compared to the β-actin with control group. Notably, the protein expression level of AMOT was significantly enriched in the β-actin&YAP1 STUPPIT group compared to the control group (Fig 2G).

To further confirm that STUPPIT can label the intermediary protein AMOT between β-actin and YAP1, we enriched the biotinylated proteins in the β-actin and YAP1 as well as the β-actin and control STUPPIT groups, and subjected the samples to mass spectrometry analysis. As expected, the intermediary protein AMOT, along with YAP1, was significantly enriched in the β-actin&YAP1 STUPPIT group compared to control (Fig 2H). These findings underscore the effectiveness of the STUPPIT tool in labeling and capturing intermediary partners between two related proteins that do not have a direct interaction.

### Validating intermediary protein labeling efficiency with Tb(N)-PupE stably expressed cells: approach II of STUPPIT

In the initial all-in-plasmids approach I of STUPPIT, we observed that the expression of 3×Flag-Tb(N)-PupE in the YAP1 group was less robust compared to the control group (Fig 2E). This discrepancy may be attributed to the inability of the IRES sequence to achieve 100% expression of the Tb(N)-PupE insert, potentially compromising the efficiency of STUPPIT (32, 33). To address this limitation, we developed approach II of STUPPIT by establishing HEK293T cells stably expressing Tb(N)-PupE (Fig 3A). This involved three constructs: PreyA-pafA-myc, the control HA-Tb(C) plasmid, and the preyC vector HA-Tb(C)-PreyC. The stable expression of 3×Flag-Tb(N)-PupE in HEK293T cells was validated through both immunofluorescence staining and immunoblotting (Fig 3B-C).

**Fig 3.**
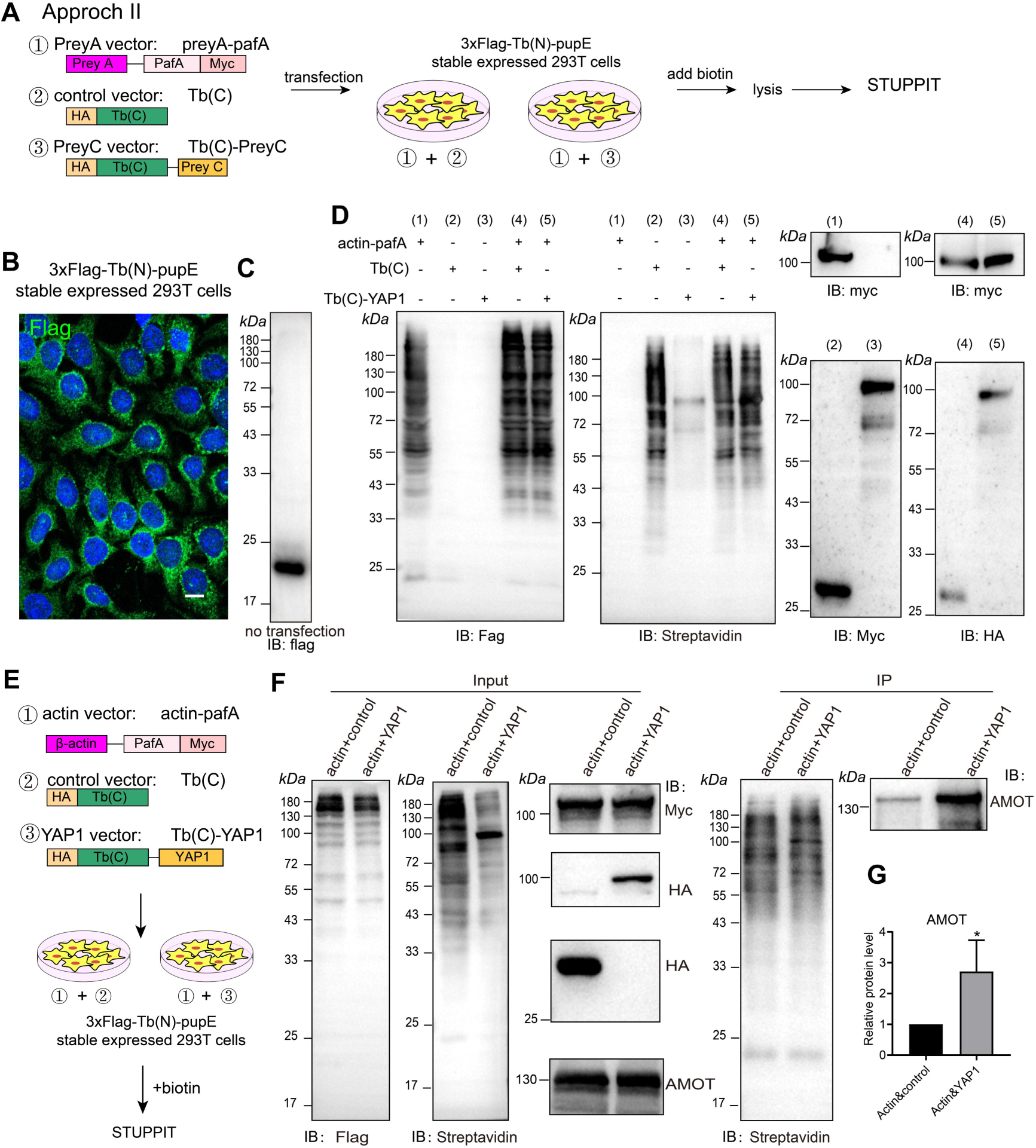
Proximity labeling of the intermediary protein AMOT between β-actin and YAP1 by approach II of STUPPIT. (A) Schematic representation of STUPPIT Approach II with HEK293T cells stably expressing 3XFLAG-Tb(N)-PupE. (B-C) Verification of stable expression of Tb(N)-PupE in HEK293T cells through immunofluorescence (B) and protein immunoblotting (C), scale bar 10 μm. (D) Immunoblotting analysis of the tagged protein under different combinations of β-actin-PafA, Tb(C)-YAP1, or Tb(C). (E) Schematic diagram illustrating the vector construction required for β-actin, control, and YAP1 according to Approach II of STUPPIT. (F) Immunoblotting validation of the captured AMOT between YAP1 and β-actin using Approach II of STUPPIT. (G) Quantification of the immunoblotting intensity of AMOT between STUPPIT of actin and YAP1 compared to the actin and control group. Data presented as mean ± SD, n=3 for each group (three independent replicates), p-value = 0.0423, two-tailed unpaired t-test.

We transfected actin-pafA-myc alone or in combination with control Tb(C)-HA or YAP-Tb(C)-HA into HEK293T cells stably expressed 3×Flag-Tb(N)-PupE. Immunoblotting analysis using the Flag antibody also revealed an array of increased bands when actin-pafA-myc was present, indicating the effectiveness of pafA in catalyzing the interaction between 3×Flag-Tb(N)-PupE and the substrate. However, transfection with actin-pafA-myc alone did not result in any bands when probed with the streptavidin antibody (Fig 3D).

We further validated the efficacy of STUPPIT approach II in labeling the intermediary protein AMOT between β-actin and YAP1 (Fig 3E). Immunoblot analysis showed comparable expression levels of FLAG-tagged proteins in both the actin-control and actin-YAP1 STUPPIT groups, indicating similar substrate expression (Fig 3F). Importantly, AMOT expression was significantly enriched in the β-actin&YAP1 group compared to the control group when utilizing approach II of STUPPIT (Fig 3F-G).

### STUPPIT Successfully Labels Other Intermediary Proteins of Known Protein Pairs

To further confirm the labeling efficiency of STUPPIT, we assessed its ability to label intermediary proteins in additional well-characterized protein pairs. The adherens junction, a mechanosensitive structure capable of sensing junctional tension, relies on α-catenin anchored to AJ by β-catenin (25, 26). This interaction allows α-catenin to engage with actin via its actin-binding domain (ABD), thereby facilitating the linkage of cell adhesion complexes to the actin cytoskeleton (Fig 4A) (34-36). We tested the β-actin/α-catenin/β-catenin axis using the STUPPIT (Fig 4B). As anticipated, both approach I and approach II of STUPPIT effectively enriched the intermediary protein α-catenin (Fig 4C-E).

**Fig 4.**
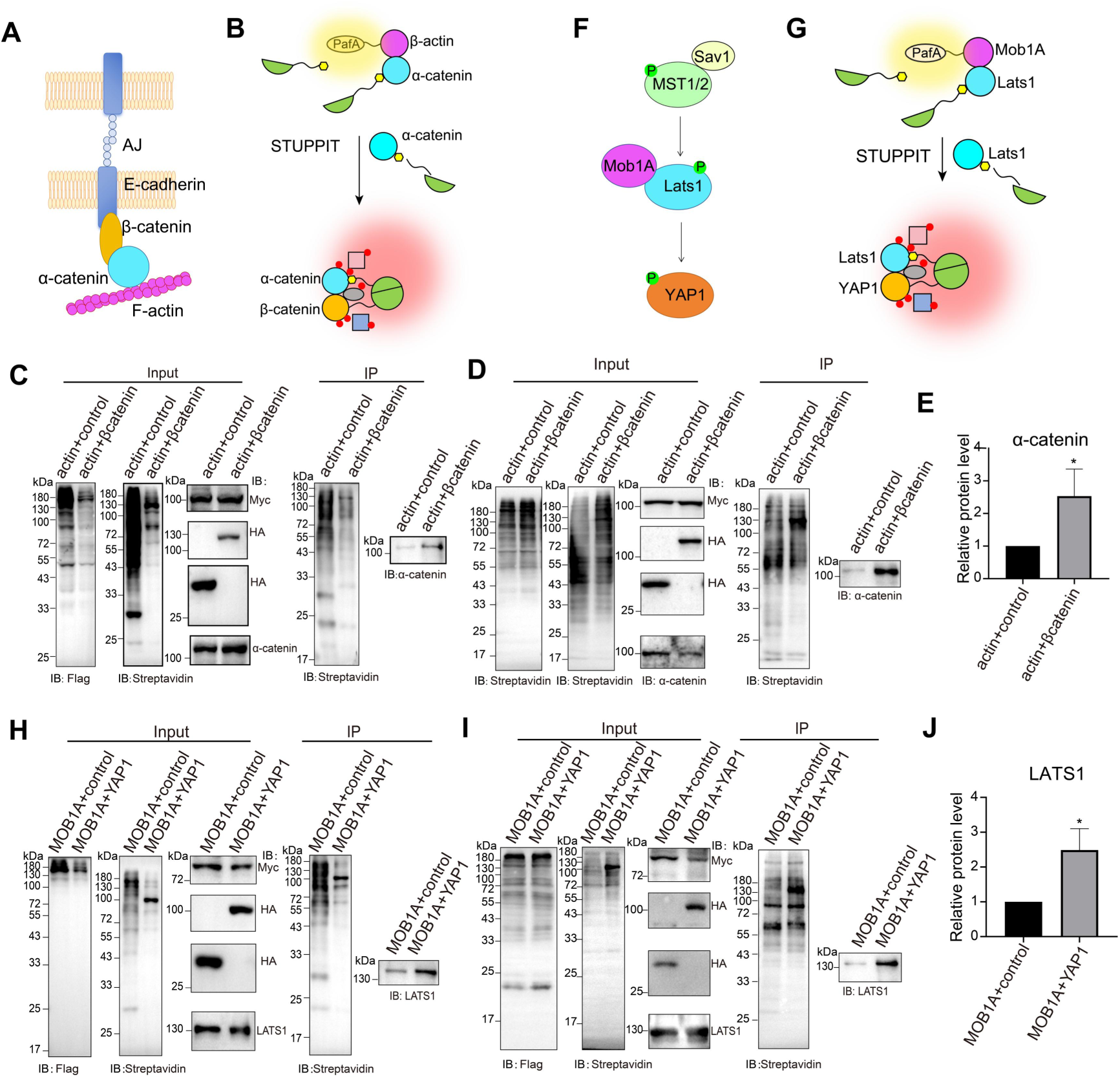
Validation of STUPPIT in capturing other known intermediary proteins. (A) α-catenin serves as the intermediary protein linking β-catenin and β-actin.(B) Schematic representation of the labeling of the intermediary protein α-catenin between β-actin and β-catenin using STUPPIT. (C) Validation of the intermediary α-catenin between β-catenin and β-actin captured by STUPPIT Approach I through immunoblotting. (D) Validation of the intermediary α-catenin between β-catenin and β-actin captured by STUPPIT Approach II through immunoblotting. (E) Quantification of the immunoblotting intensity of α-catenin between the STUPPIT of β-actin&β-catenin compared to the β-actin&control group. Data shown as mean ± SD, n=3 for each group (three independent replicates), p-value = 0.0330, two-tailed unpaired t-test. (F) Description of the intermediary protein LATS1 positioned between MOB1A and YAP1. (G) Schematic representation of the labeling of the intermediary protein LATS1 between MOB1A and YAP1 using STUPPIT. (H) Validation of the captured LATS1 between MOB1A and YAP1 by STUPPIT Approach I through immunoblotting. (I) Validation of the captured LATS1 between MOB1A and YAP1 by STUPPIT Approach II through immunoblotting. (J) Quantification of the immunoblotting intensity of LATS1 between the STUPPIT of MOB1A&YAP1 compared to the MOB1A&control group. Data presented as mean ± SD, n=3 for each group (three independent replicates), p-value = 0.0136, two-tailed unpaired t-test.

Furthermore, we also extended the application of STUPPIT to the MOB1A-LATS1-YAP1 cassette, which represents typical components of the Hippo pathway (Fig 4F) (27-29). LATS1 serves as the intermediary protein between Mob1A and YAP1 (Fig 4G). As expected, both approach I and approach II of STUPPIT successfully enriched LATS1 (Fig 4H-J). These results, combined with the enrichment of AMOT between β-actin and YAP1 by two approaches, further validate of the ability of STUPPIT in labeling intermediary proteins.

### Applying STUPPIT to Identify Novel Intermediary Proteins of SMAD4 and β-catenin, Key Components of BMP and Wnt Pathways

Finally, we applied STUPPIT to identify novel intermediary proteins between key components of the BMP and Wnt pathways. Reciprocal interactions between these pathways are critical to control stem cell behaviors during tissue development and homeostasis, such as the small intestine (30) and hair follicles (31). These pathways are known to counteract each other, potentially through competition for common binding partners (Fig 5A).

**Fig 5.**
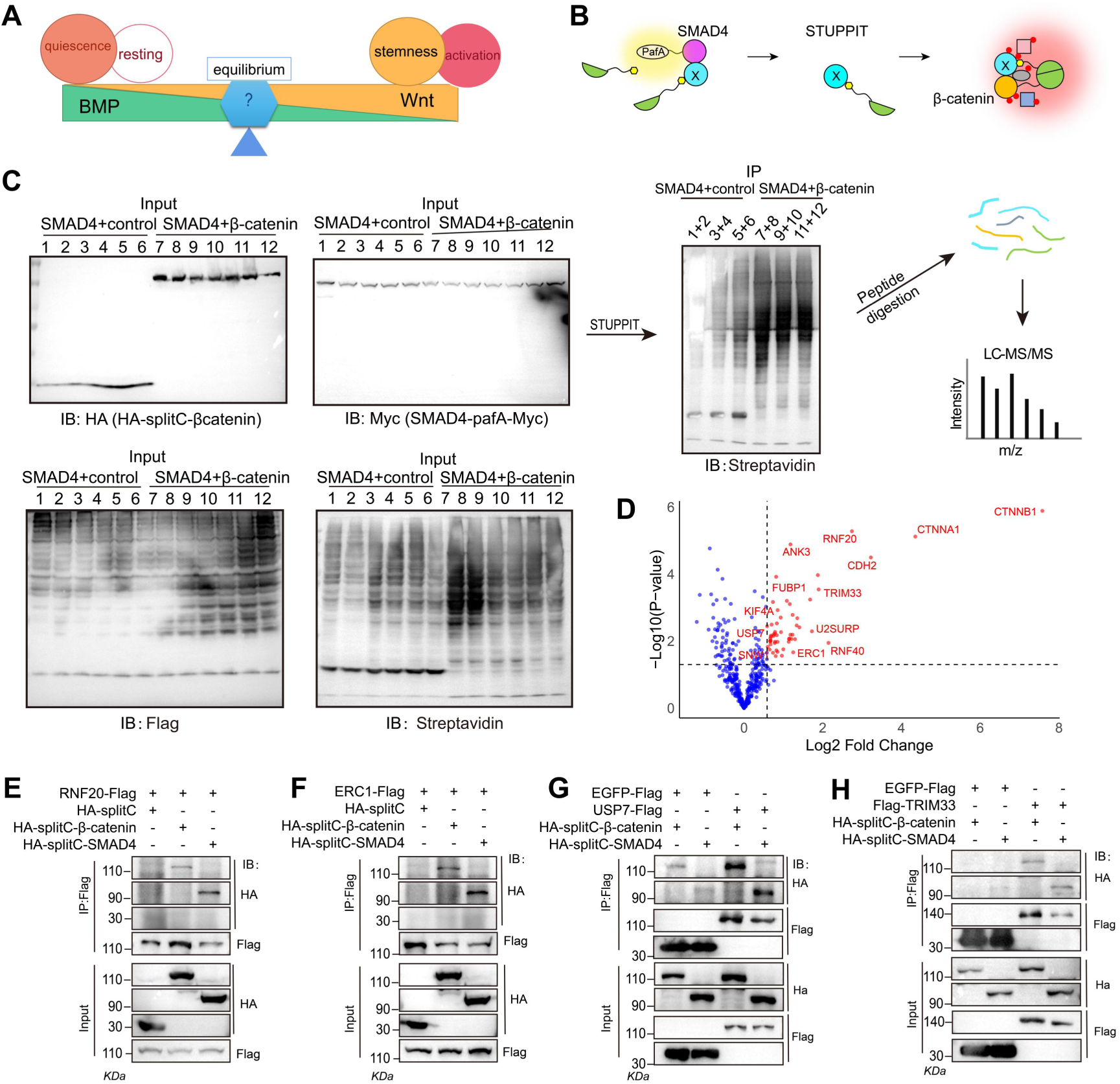
Application of STUPPIT in capturing novel intermediary proteins between SMAD4 and β-catenin, core components of BMP and Wnt pathway. (A) Diagram dictating the counteract role of BMP and Wnt signaling in controlling stem cell behaviors. (B) Diagram of labeling the intermediary proteins between SMAD4 and β-catenin using STUPPIT. (C) STUPPIT labeled intermediary proteins between SMAD4 and β-catenin were enriched and analyzed by mass spectrometry. (D). Volcano plots showing possible intermediary proteins between SMAD4 and β-catenin. Significantly enriched proteins with 1.5 fold change are labeled as red dots. Proteins with unique peptides>9 in the SMAD4&β-catenin group were selected. (E-H). Immunoprecipitation assays showing interactions between RNF20 (E), ERC1 (F), USP7 (G), TRIM33 (H) with SMAD4 and β-catenin, respectively.

We selected β-catenin and SMAD4 as prey proteins for STUPPIT (Fig 5B). Biotinylated proteins labeled by STUPPIT were enriched and subjected to mass spectrometry analysis (Fig 5C). The proteomics data implied that proteins such as RNF20, ERC1, RNF40, TRIM33, USP7 might be the intermediary proteins between β-catenin and SMAD4 (Fig 5D and S2 Fig). Notably, TRIM33 has been previously reported to interact with β-catenin (37) and SMAD4 (38), separately. We confirmed the interaction of RNF20, ERC1, USP7, and TRIM33 with β-catenin and SMAD4 through immunoprecipitation (Fig 5E-H and S3 Fig). Together, these data validated the ability of STUPPIT to label and capture the intermediary proteins between two non-interacting proteins, thereby facilitating the investigation of signaling transduction and crosstalk.

## Discussion

Identifying novel protein-protein interactions is key to understand signal transduction and crosstalk in cells and tissues. Our study introduces a novel tool, STUPPIT, which addresses a critical gap in proximity labeling by enabling the direct labeling and capture of intermediary proteins between non-interacting partners. This advancement significantly enhances our ability to dissect complex signaling pathways and their underlying mechanisms.

Proximity labeling tools such as TurboID and APEX have been instrumental in tagging endogenous proteins that interact with a specific bait protein, allowing their enrichment and identification through mass spectrometry. While split-TurboID methods can detect proximal partners of interacting proteins, they rely on close proximity to ensure intact TurboID activity. However, these methods are limited in their ability to directly label and capture endogenous intermediary proteins between associated but non-interacting proteins. STUPPIT overcomes this limitation by not requiring direct interaction between the two bait proteins, making it a powerful tool for studying signaling pathways where key components may not directly interact.

It is worth noting that other split-proximity labeling technologies such as split-APEX2 (39) or split-HaloTag system (40) could theoretically replace split-TurboID in the STUPPIT system for labeling intermediary proteins in cells. However, the length of the split-fragment linked to Pup(E) must be carefully optimized to ensure efficient tagging of the modified Pup(E) to the lysine amino acid in the intermediary proteins by pafA. This optimization is critical for maintaining the specificity and efficiency of the labeling process. Moreover, STUPPIT not only labels intermediary proteins and prey C but also identifies proximity proteins between the intermediary protein and prey C. This dual functionality is advantageous, as it further enriches the core components of intermediary proteins and prey C proteins, facilitating subsequent investigations into their relationship. This also implies that STUPPIT can be used as an alternative to split-TurboID for labeling proximal proteins between two interacting proteins. The successful identification of intermediary proteins between SMAD4 and β-catenin, key components of the BMP and Wnt pathways, highlights the broad application of STUPPIT in exploring the crosstalk between signaling pathways. These intermediary proteins, such as TRIM33, interact both with SMAD4 and β-catenin, implying potential competitive binding between these pathways. This finding underscores the potential of STUPPIT to reveal novel regulatory mechanisms in signaling networks.

One limitation shared by STUPPIT, similar to split-TurboID, is the presence of background noise in control group. To mitigate this issue, it is advisable to maintain only one variable in the STUPPIT system between two prey proteins. When identifying true intermediary proteins following STUPPIT and mass spectrometry analysis, it is essential to account for the background labeling in the control group, even if the protein is enriched in the experimental group. Such analyses should be taken into consideration to narrow down potential candidates. Subsequent validation steps, such as immunoprecipitation, are necessary to confirm the true intermediary proteins.

Future work should focus on further optimizing the STUPPIT system to reduce background noise and improve labeling efficiency. Additionally, exploring the application of STUPPIT in different cell types and tissues will provide broader insights into the dynamic nature of protein-protein interactions in various biological contexts. The identification of novel intermediary proteins using STUPPIT could lead to new therapeutic targets and a deeper understanding of signaling pathways in health and disease.

## Materials and Methods

### Lead contact

Any requests for further information and resources should be directed to Wenxiu Ning, the lead contact on this paper.

### Materials availability

This study generated new plasmids were available from Dr. Ning.

### Experimental model and subject details Cells

All cells were cultured with a DMEM medium (Thermo Fisher, #12634-010, US) containing 10% fetal bovine serum (Vivacell#C04001-500, China) and 1% penicillin-streptomycin (Beyotime, China) at 37 °C with 5% CO2. HEK293T cells were used for immunoprecipitation assays. Hela cells were used for immunofluorescence staining. DNA transfection of HEK293T and Hela cells was performed by the HighGene plus transfection reagent (ABclonal, China). To establish 3XFag-Tb(N)-Pup(E) stably expressed HEK293T cells, psPAX2, pMD2G, pPHAGE-EF1A-puro-3×Flag-Tb(N)-Pup(E) plasmids and HEK293T cells were used for virus generation with PEI reagent (Polysciences#24765, US). The produced packaging lentiviruses were then added to the HEK293T cells with 10 μg/mL polybrene (Biosharp#BL628A, China), and incubated for 48 hours, after which cells were selected with 1 μg/mL puromycin for 3 days.

### Plasmids construction

PafA-Myc DNA fragment and Pup(E) peptide were amplified from plasmids pEF6a_kozak_CD28_PafA-Myc and pEF6a-HB-Pup(E) provided by Prof. Min Zhuang (Shanghai Tech University). IRES DNA fragment was amplified from plasmid pLVX-mgSrtA-IRES-ZsGreen provided by Prof. Peng Chen (Peking University). Tb(N) and Tb(C) DNA fragments (split at the amino acid site L73/G74) were amplified from a TurboID plasmid pLVX-1xFlag-TurboID provided by Prof. Wenxiang Fu (Yunnan University). YAP1, β-actin, MOB1A, β-catenin, RNF20, ERC1, USP7 and TRIM33 were amplified from cDNA of human HaCaT keratinocytes respectively. pCMV-Tag2B plasmid was provided by Prof. Jianwei Sun. pHAGE-EF1a-puro was provided by Prof. Maorong Chen from Yunnan University. Self-made plasmids including pCMV-actin-pafA-Myc, pCMV-HA-Tb(C)-YAP1, pCMV-HA-Tb(C), pCMV-HA- Tb(C)-IRES-3×Flag-Tb(N)-Pup(E), pCMV-HA-Tb(C)-IRES-3×Flag-Tb(N)-Pup(E), pPHAGE-EF1A-puro-3×Flag-Tb(N)-Pup(E) are available upon request from the Lead Contact.

Infusion cloning primers were designed using Vazyme’s software, CE Design (available on https://www.vazyme.com). Prepare Linearized Vectors by digesting the circular vector with relative restriction endonucleases. Prepare inserts by PCR amplification using high-fidelity polymerase. Calculate vectors and inserts input (ratio of vector to insert is about 1:2) after determination of their concentration. Recombine the linearized vector, insert and Exnase II (ClonExpress II One Step Cloning Kit, C112-01) following the description. Incubate at 37°C for 30 min, then chill on ice. Transform the recombination products to the competent cells. Identification of recombinant products were performed by colony PCR with primers designed. Correct plasmid or colony was selected and sent for DNA sequencing.

### Cell transfection

Cells were seeded at 3x10^6^ cells per 6 cm plates to achive approximately 50% confluence. The following day, transfection was performed following protocols provided for HighGene reagents (Abclonal, China). First, the culture medium was replaced with fresh DMEM. In a centrifuge tube, 400 μL of DMEM medium was added, followed by 4 μg of plasmid DNA, and then 8 μL of HighGene transfection reagent was added. After gently vortexing to mix the solution, thoroughly, it was incubated at room temperature for 15 minutes. The transfection mixture was then evenly and gently added to each dish, and the dish was gently shaken to ensure even distribution of the mixture over the cells. Finally, the dishes were placed in the incubator for cell culture, allowing the cells to uptake the exogenous DNA.

### Cell lysis

Gently wash the cells twice with PBS, being careful to avoid dislodging surrounding cells. Aspirate the PBS and add an appropriate volume of RIPA lysis buffer (400 μL for a 6 cm dish, 1 mL for a 10 cm dish). Use a cell scraper to gently scrape the cells from the culture dish and allow the lysate sit for 10 minutes to ensure complete lysis. Then transfer the lysate to a 1.5 mL centrifuge tube and centrifuge at 13,000 g for 10 minutes. After centrifugation, transfer the supernatant to a new 1.5 mL centrifuge tube and add PMSF at a ratio of 1:200. Aliquot a portion of the lysate and mix it with 3x sample buffer for protein analysis before ultrafiltration. The remaining lysate will be used for ultrafiltration processing. The RIPA buffer (100 mL) was prepared as follows: 3 mL 5M NaCl, 1 mL 0.5 M EDTA, pH 8.0, 5 mL 1M Tris-HCl, pH 8.0, 1 mL NP-40, 5 mL 10% sodium deoxycholate, 10% SDS: 1 mL, and ddH_2_O to bring the volume to 100 mL.

### Ultrafiltration to remove free biotin

Add 2 mL of pre-chilled PBS to the prepared ultrafiltration column (Pall Microsep MCP003C4-5 mL), and add 20 μL of PMSF (final volume of the ultrafiltrate is 4 mL, with a PMSF solution ratio of 1:200). Add the remaining lysate to the ultrafiltration column and bring the total volume to 4 mL. Centrifuge the column at 4000 g for 1 to 1.5 hours. Repeat this step three times to ensure complete removal of free biotin (Biotin, Sigma#B4501). The final volume of the lysates after the ultrafiltrate is about 400 μL after the final centrifugation. Transfer the ultrafiltrate to a new 1.5 mL centrifuge tube. Aliquot 80 μL of the lysate and add 40 μL of 3x sample buffer for input. The remaining lysate will be used for streptavidin affinity immunoprecipitation. If not processed immediately, add PMSF at a ratio of 1:100 and store at -80°C.

### Streptavidin affinity immunoprecipitation

Gently resuspend the Streptavidin Agarose Resin (YEASEN#20512ES08, China) using a pipette to evenly disperse the beads in the solution. Transfer the required volume of beads into a 1.5 mL centrifuge tube on ice (15 μL for a 6 cm tube, and 30 μL for a 10 cm tube). Centrifuge at 4°C, 3000 rpm, for 2 minutes, and remove the supernatant. Add 1 mL of 1xTBS by gently pipetting, centrifuging, and discarding the supernatant. Repeat this washing step three times. Add an appropriate volume of PBS buffer, gently agitate to re-suspend the beads, and distribute evenly into each sample. Add PBS to bring the volume of each 1.5 mL centrifuge tube to 1ml. Seal the samples and incubate at room temperature on a rocking platform for 1 hour. Centrifuge and remove the supernatant. Wash the beads with 1 mL PBS or PBST buffer containing 0.05% Tween, then centrifuge to wash the beads 3-4 times, and remove the supernatant. Add an appropriate amount of 1x sample buffer (typically 65 – 75 μL), heat at 95°C for 10 minutes, and then freeze the samples at -20°C. The 3x Sample Buffer (85 mL) was made as follow: 18.8 mL 1M Tris pH 6.8, 10.0 mL 20% SDS, 30.0 mL glycerol, 10.0 mL 0.4% bromophenol blue solution, 16.2 mL H_2_O. Add β-mercaptoethanol or 15 mM DTT to individual aliquots when using (850 μL 3x sample buffer +150 μL BME).

### Immunoprecipitation of Flag fusion proteins

Transfect HEK293T cells with plasmids expressing Flag-fused proteins. After 48 hours, gently wash the cells with PBS. Lyse the cells by adding 500 μL of RIPA buffer (containing PMSF, and protease inhibitor) on ice. Mix thoroughly and transfer the lysate to a 1.5 mL centrifuge tube. Centrifuge at 12000 rpm for 5 minutes at 4°C. Collect the supernatant and reserve 100 μL as input. Gently resuspend the anti-Flag affinity beads (Beyotime#P2282, China) in TBS. Transfer the required volume of beads to a clean centrifuge tube (8 μL for a 6 cm culture dish). Add 500 μL of 1xTBS to gently suspend the beads. Centrifuge at 6,000g for 30 sec at 4°C and discard the supernatant. Repeat this step twice. Resuspend the beads in 20 μL of 1xTBS per sample. Add about 20 μL of bead suspension to each sample of cell lysate. Incubate the samples at 4°C for 1-2 hours or overnight. After incubation, centrifuge at 6,000g for 30 sec at 4°C. Discard the supernatant without disturbing the beads. Wash the bead by adding 500 μL of 1xTBS, gently suspending and centrifuging at 6,000 g for 30 sec at 4°C. Discard the supernatant and repeat this washing step three times. Finally, dissolve in 60 μL of 1x sample buffer and heat at 95°C for 5 minutes.

### Western blotting

Samples were separated by SDS-PAGE and transferred onto PVDF membranes, then blocked with 5% nonfat milk at room temperature for 1 hour. Then membranes were incubated with primary antibody diluted in 2% nonfat milk for 2.5 hours at RT, followed by washing with 0.1% Tween-20 in PBS three times. Subsequently, the membranes were incubated with HRP-conjugated secondary antibody diluted with 2% nonfat milk for 1 hour at RT. Finally, the samples were washed again with 0.1% Tween-20 in PBS and imaged using chemiluminescence. Antibodies used were listed as follows: Horse Anti-mouse IgG, HRP-linked Antibody, CST#7076; Goat Anti-rabbit IgG, HRP-linked Antibody, CST#7074; HRP-labeled Streptavidin, Beyotime#A0303; Rb anti Alpha E-Catenin, proteintech#12831-1-AP; Rb anti-AMOT, proteintech#24550-1-AP; ms anti-Flag M2, Sigma#F1804; ms anti-HA tag, Invitrogen#26183; Rb anti-LATS1, proteintech#17409-1-AP; ms anti-myc, abclonal#AE010. Original uncropped images of western blotting were included in S1 Raw images in the supplementary information.

### Immunofluorescence staining

Cells were plated in 24 well plates containing glass-coverslips at a density of 1x10^5^ cells per well. Transfect the related plasmids into the wells. After 48 hours, remove the medium and fix the cells with 4% paraformaldehyde for 10 min. Wash the wells three times with PBS containing 0.1% Triton X-100 for 5 min each time. Block with 3% BSA buffer for 30 min. Then incubate with 25 μL of diluted primary antibodies for 1 hour. Wash the wells three times with PBS containing 0.1% Triton X-100 for 5 min each time. Next, incubate with 25 μL of diluted secondary antibodies and DAPI for 1 hour. Wash the wells three times with PBS containing 0.1% Triton X-100 for 5 min each time. Finally, mount the coverslips on slides with an anti-fade solution.

### Imaging

Slides were imaged using an inverted Zeiss LSM800 confocal microscope with Airyscan using 63x oil-immersion objective lens and processed using Fiji image.

### Mass spectrometry sample preparation and data analysis

First, STUPPIT streptavidin affinity precipitation was conducted to purify the biotinylated protein samples. Subsequently, the purified biotinylated protein samples were first run in a 10% SDS-PAGE gel, followed by excision of the protein bands of interest. The excised bands were then submitted to the mass Mass Spectrometry Facility of Yunnan University for mass spectrometry analysis. In-gel digestion and LC-MS/MS analysis were performed following the protocol as described (41). Protein identification and ion intensity quantification were performed by MaxQuant software (version 1.6.4.0) (42). The protein intensity is the sum of all peptides identified from this protein. Candidate proteins were selected based on the log2intensity and numbers of unique peptide that were enriched in the experimental group compared to control. NaN is a missing value in the data. This means that none of the peptides from these proteins were detected by mass spectrometry, which may be due to extremely low peptide abundance. Proteins identified by mass spectrometry in three biological replicates following STUPPIT between actin and YAP1, or SMAD4&β-catenin were included in S1 Data.

### Quantifications and statistical analysis

Prior to performing statistical tests, Normality of data distribution was assessed by D’Agostino–Pearson test using GraphPad Prism 5 software. Data are represented as the mean ± SEM, and were judged to be statistically significant when p value < 0.05 by unpaired Student’s t test (ns = not significant, *, p < 0.05; **, p < 0.01, ***, p < 0.001, ****, p < 0.0001). All data were performed at least three independent replicates. All images were analyzed using FIJI ImageJ. Statistical analysis was performed using GraphPad Prism 5 software and Microsoft Excel.

## Author Contribution

Lin Xie, Hua Li, Lijuan Gao: Investigation, Methodology, Validation. Lin Xie: Formal analysis, validation. Gangyun Wu: Validation. Hua Li: Conceptualization, Writing – review & editing, Supervision, Funding acquisition. Wenxiu Ning: Conceptualization, Formal analysis, Writing – original draft, Writing – review & editing, Supervision, Resources, Funding acquisition. Lin Xie, Hua Li, Lijuan Gao contributed equally to this work.

## Declaration of interests

The authors declare no competing interests

## Acknowledgements

We are grateful to Professor Xuna Wu for mass spectrometry analysis, the core facilities of Yunnan University for imaging, Professor Wenxiang Fu, Maorong Chen, Jianwei Sun, Min Zhuang and Peng Chen for kindly gifts of plasmids as indicated in methods. This study was supported by the National Natural Science Foundation of China No. 32270846, Wenxiu Ning; and Applied Basic Research Foundation of Yunnan Province No. 202401AT070443, Hua Li..

## Supporting information

**S1 Fig.**
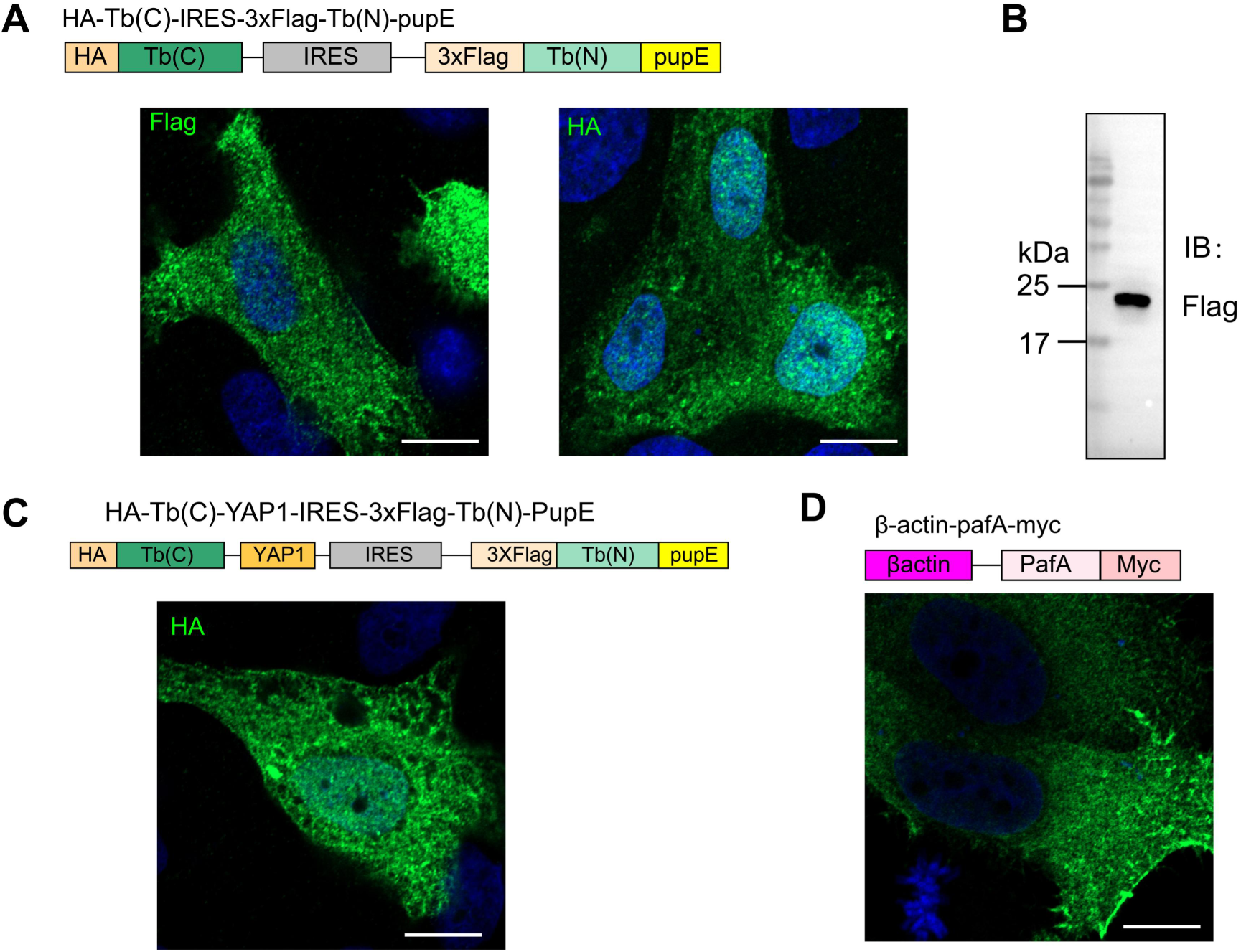
Validation for the expression of the YAP1 and actin related plasmids. (A) Immunofluorescence staining of Flag and HA tag in control vector. (B) Immunoblotting of 3xFlag-Tb(N)-PupE in STUPPIT when expressed alone in HEK293T cells. (C) Immunofluorescence staining of HA tag in YAP1 vector. (D) Immunofluorescence staining of Myc tag in actin-pafA-Myc vector. Scale bars are all 10 μm.

**S2 Fig.**
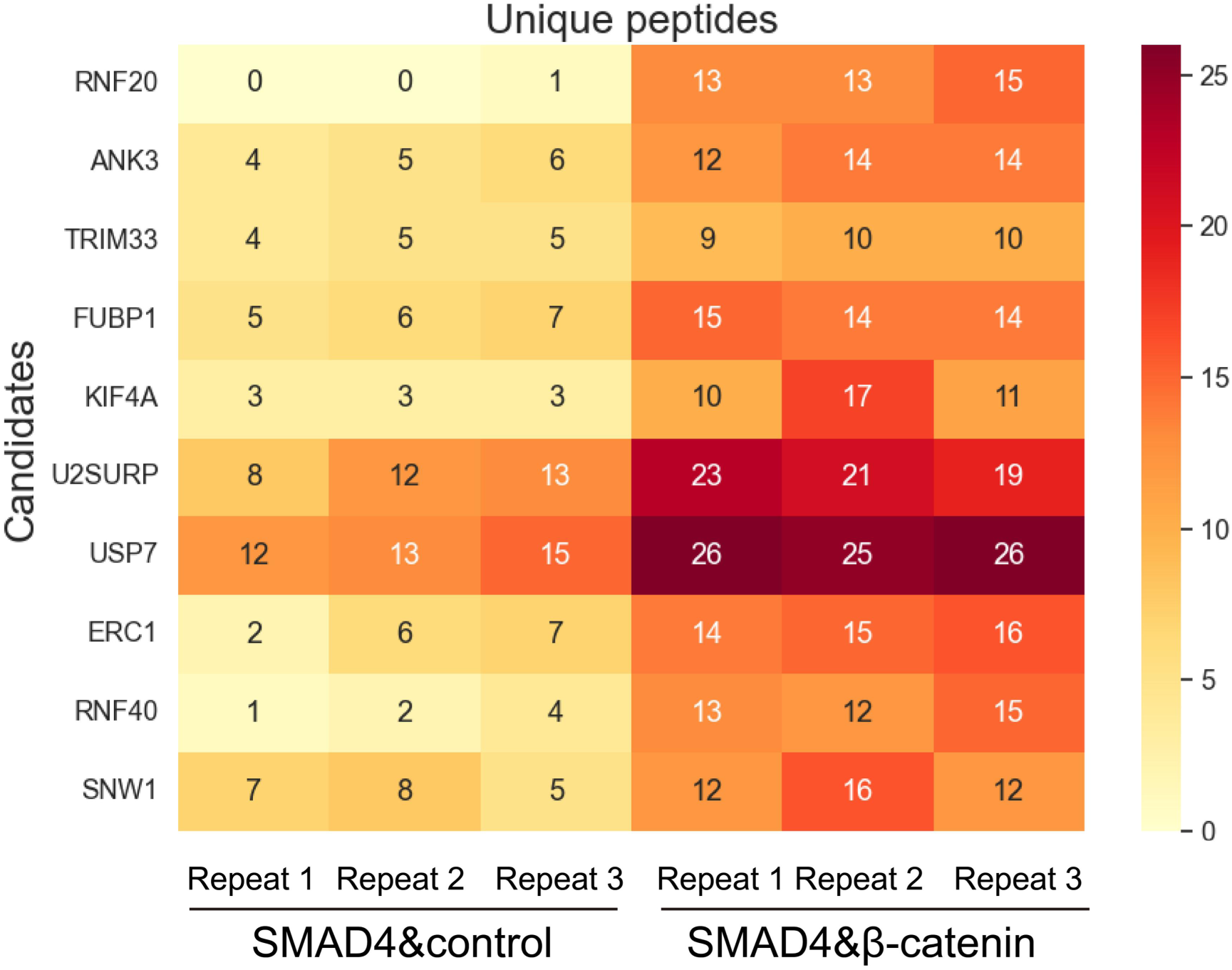
Heatmap showing the number of the unique peptide of the indicated candidates in SMAD4&β-catenin and SMAD4&control groups identified by mass spectrometry analysis.

**S3 Fig.**
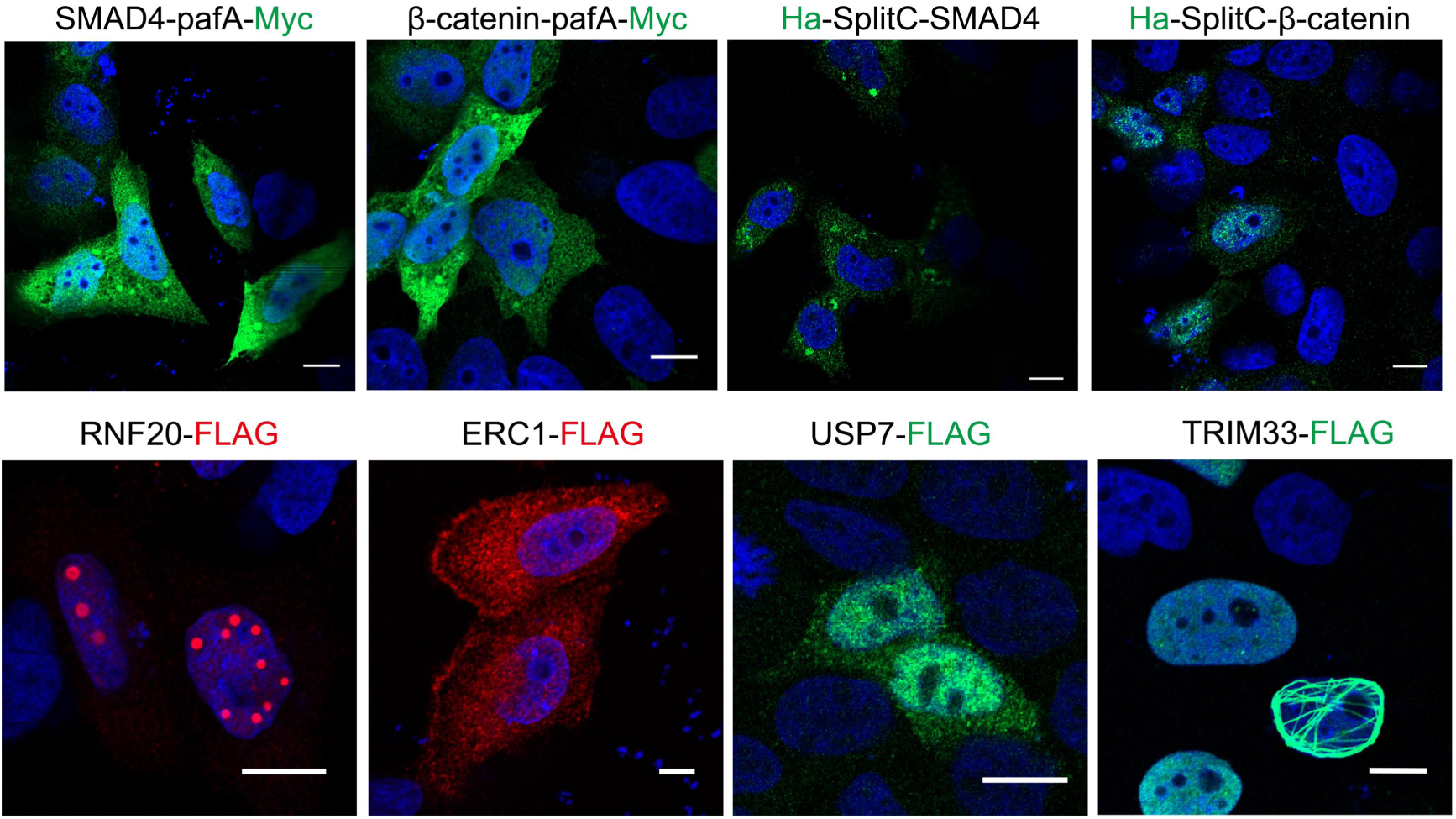
**Validation for the expression of the indicated proteins by immunofluorescence staining.**

**S1 Raw images. PDF file containing un-cropped images of all western blots in this manuscript.**

**S1 Data. Proteins identified by mass spectrometry in three biological replicates following STUPPIT enrichment for intermediary proteins between actin and YAP1, or SMAD4&βcatenin, respectively.**

## Notes

### Competing Interest Statement

The authors have declared no competing interest.

